# Soluble Urokinase Plasminogen Activator Receptor Primes Macrophages and Worsens Heart Failure with Preserved Ejection Fraction

**DOI:** 10.64898/2026.06.04.730230

**Authors:** Anand Prakash Singh, Parisa Shabani, Anis Ismail, Rajesh Chaudhary, Afnan Alzamrooni, Tahra Luther, Matthew Nho, Rachel Lopez, Chetna Soni, Sascha N. Goonewardena, Salim Hayek, Ahmed Abdel-Latif

**Affiliations:** Division of Cardiology, Department of Internal Medicine, Frankel Cardiovascular Center, University of Michigan, Ann Arbor, MI 48105, USA; Ann Arbor VA Healthcare System, 2215 Fuller Rd, Ann Arbor, MI 48105, USA; University of Texas Medical Branch, Department of Internal Medicine, Galveston, TX, USA; Department of Internal Medicine, Division of Infectious Diseases, University of Michigan, Ann Arbor, MI 48109, USA

## Abstract

**Background:** Heart failure with preserved ejection fraction (HFpEF) is a systemic inflammatory syndrome with few effective therapies. Soluble urokinase plasminogen activator receptor (suPAR), a circulating immune-derived glycoprotein, independently predicts adverse outcomes in HFpEF beyond natriuretic peptides, but whether it is a causal driver or a passive marker of inflammatory burden has remained unresolved.

**Methods:** We tested the hypothesis that elevated circulating suPAR is sufficient to amplify HFpEF by acting on the innate immune system. suPAR-transgenic (suPAR-Tg) and wild-type mice were subjected to a cardiometabolic two-hit model (high-fat diet plus L-NAME) for 15 weeks. Cardiac structure and diastolic function were assessed by serial echocardiography alongside blood pressure, glucose tolerance, and gravimetric endpoints, and left ventricular tissue was profiled by bulk RNA sequencing with in silico cellular deconvolution. Myeloid populations in the heart, spleen, and peripheral blood were quantified by spectral flow cytometry and corroborated by galectin-3 immunofluorescence, and the direct effect of suPAR on macrophages was tested by priming bone marrow–derived macrophages with recombinant suPAR before LPS and IFN-γ stimulation.

**Results:** Sustained suPAR elevation worsened the established HFpEF phenotype, producing greater diastolic dysfunction (higher E/e′ and E/A ratios) and pulmonary congestion without altering blood pressure or ejection fraction, indicating a mechanism downstream of the canonical hemodynamic stimulus. Bulk RNA sequencing of left ventricular tissue revealed a coordinated transcriptional shift, with suppression of mitochondrial oxidative phosphorylation and amplification of innate and adaptive immune programs, including interleukin-1β production, leukocyte chemotaxis, and antigen presentation. Spectral flow cytometry demonstrated stepwise expansion of CCR2⁺ inflammatory monocytes and macrophages across cardiac, splenic, and peripheral compartments, corroborated in situ by increased galectin-3⁺ macrophage density. In vitro, recombinant suPAR was not a stand-alone inflammatory ligand but instead primed bone marrow–derived macrophages to markedly amplify TNF-α, IL-1β, IL-6, and NLRP3 responses to LPS and IFN-γ.

**Conclusions:** Together, these findings establish that elevated suPAR is sufficient to act as an upstream amplifier of HFpEF, identify the CCR2⁺ inflammatory monocyte-macrophage axis as its proximate effector, and convert two decades of epidemiologic association into a mechanistically grounded, therapeutically tractable hypothesis with immediate relevance to clinical-stage anti-suPAR antibodies.

## Introduction

Heart failure with preserved ejection fraction (HFpEF) accounts for more than half of all heart failure cases worldwide and carries a 5-year survival of approximately 24% following an index hospitalization, outcomes comparable to those of many advanced malignancies (Redfield & Borlaug, 2023; Borlaug et al., 2024). Only sodium-glucose cotransporter 2 inhibitors reduce heart failure hospitalization in HFpEF (Anker et al., 2021; Solomon et al., 2022), and no therapy has matched the mortality benefit of guideline-directed therapy in heart failure with reduced ejection fraction. The therapeutic shortfall reflects, in large part, our incomplete grasp of the disease’s mechanistic architecture.

The contemporary paradigm frames HFpEF as a systemic inflammatory syndrome in which extracardiac comorbidities (obesity, type 2 diabetes, chronic kidney disease, hypertension) generate sustained low-grade inflammation that, through coronary microvascular endothelial dysfunction and oxidative stress, recruits monocyte-derived macrophages to the myocardium and drives fibrosis and diastolic stiffening (Paulus & Tschöpe, 2013; Paulus & Zile, 2021). This framework is supported by multiple converging lines of evidence: elevated circulating inflammatory cytokines in HFpEF patients, NLRP3 inflammasome activation in the cardiometabolic two-hit mouse model (Schiattarella et al., 2019; Schiattarella et al., 2021), and the recent demonstration that CCR2⁺ monocyte-derived macrophages drive cardiac hypertrophy, fibrosis, and diastolic dysfunction in early HFpEF and pressure-overload models (Patel et al., 2018; Tucker et al., 2024; Raman et al., 2025). Yet the upstream circulating signals that selectively expand and activate this inflammatory pathway, and that might therefore offer tractable therapeutic entry points, remain incompletely defined.

Soluble urokinase plasminogen activator receptor (suPAR), a circulating signaling glycoprotein produced predominantly by activated immune cells, has emerged as one of the most consistent prognostic biomarkers across cardiovascular and renal disease (Eugen-Olsen et al., 2010; Reiser, 2026). In heart failure populations, baseline suPAR independently predicts mortality and hospitalization with risk-prediction improvement beyond natriuretic peptides (Hayek et al., 2023). In a recent ancillary analysis of the TOPCAT trial, patients with HFpEF and baseline suPAR in the highest tertile suffered more than a twofold increased risk of cardiovascular death, cardiac arrest, or heart failure hospitalization compared with patients in the lowest tertile, with suPAR providing a 0.123-point improvement in C-statistic beyond age, sex, race, and natriuretic peptides (Hutten et al., 2025). Causality, however, has not been established. suPAR could reflect cumulative inflammatory burden, serve as a surrogate for the comorbidities that drive HFpEF, or actively participate in the inflammatory cascade; these possibilities have sharply different therapeutic implications.

A causal molecular framework for suPAR’s pathological effects exists in other organs. In podocytes, suPAR engages the αvβ3 integrin in a tripartite complex with APOL1 to drive proteinuric kidney disease (Wei et al., 2008; Hayek et al., 2017); in atherosclerosis, suPAR-transgenic mice exhibit accelerated lesion development through monocyte priming and enhanced chemotaxis (Hindy et al., 2022). Whether the same upstream-amplifier logic applies to cardiac biology in HFpEF, and whether soluble uPAR engages cardiac and peripheral macrophages to potentiate the inflammatory cascade that defines the disease, have not been directly tested. This question has immediate translational relevance: WAL0921, a humanized anti-suPAR monoclonal antibody, has entered Phase 2 clinical trials for chronic kidney diseases (Walden Biosciences, 2025).

To address this gap, we used a genetically defined suPAR-transgenic mouse line subjected to an established cardiometabolic HFpEF model (high-fat diet plus L-NAME). Sustained elevation of circulating suPAR worsened the established HFpEF phenotype, drove a coordinated transcriptional shift toward innate and adaptive immune activation in the myocardium, and expanded CCR2⁺ inflammatory monocytes and macrophages across cardiac, splenic, and peripheral compartments. *In vitro,* recombinant suPAR primed bone marrow–derived macrophages for amplified inflammatory cytokine and NLRP3 inflammasome responses to LPS and IFN-γ. Together, these findings establish that elevated suPAR is sufficient to act as an upstream amplifier of HFpEF, identify the CCR2⁺ inflammatory monocyte-macrophage axis as its proximate effector, and convert two decades of epidemiologic association between suPAR and adverse heart failure outcomes into a mechanistically grounded and therapeutically tractable hypothesis.

## Methods

### Animals

All animal experiments were performed in accordance with the approved University of Michigan Institutional Animal Care and Use Committee (IACUC) protocol. Sex-specific differences in responses to cardiometabolic syndrome are known to exist in both mice and humans^27,28^. Since female mice are less susceptible to developing cardiac phenotype than males^29^, we used 8-10-week-old male C57BL/6 mice (Jackson Laboratory, Bar Harbor, ME) and suPAR transgenic mice (suPAR-Tg) for this study. Baseline measurements of body weight and echocardiogram were taken before they were randomized to receive a low-fat diet (LFD, #D12450J, Research Diet, USA) or a high-fat diet (HFD, #D12492, Research Diet, USA) supplemented with nitric oxide synthase inhibitor, Nω-nitro-L-arginine methyl ester (L-NAME, 0.75 g/L) (#N5751, Sigma, USA) in drinking water. All mice were housed at an ambient temperature of 23 °C, at a density of 2-3 mice/cage in 12 h of light:12 h of darkness with lights off at 07:00 (Zeitgeber, ZT 0) and lights on at ZT 12, and had ad libitum access to food and water throughout the 15-week study duration. Food, water, and cages were changed twice weekly for all mice.

### Intraperitoneal glucose tolerance test (IPGTT)

An intraperitoneal glucose tolerance test (IPGTT) was performed at weeks 5 and 15 after 5 hours of fasting, using 2 g/kg body weight (3-6 mice/group). Blood glucose readings were measured at 0, 15, 30, 60, 90, and 120 minutes after the bolus dose via tail-vein prick using Accu-Chek® Performa II (Roche, USA). The area under the curve (AUC) was calculated using the trapezoidal rule^30^.

### Blood pressure measurements

Blood pressure measurements were conducted using a CODA Monitor (Kent Scientific) via the tail-cuff method, according to the manufacturer’s instructions, after the mice were acclimatized and settled in a quiet procedure room with no disturbance. Briefly, an occlusion (O) tail cuff and a volume pressure recording (VPR) cuff were placed on the tail of the mouse, and the O-cuff was inflated to impede blood flow to the tail before the cuff was deflated slowly, and the return blood flow was measured by using a VPR sensor. Body temperature was measured at the base of the tail using a laser thermometer. All measurements were recorded in non-anesthetized mice, and body temperature was maintained using a thermal pad.

### Echocardiography measurements

All echocardiographic measurements were recorded in anesthetized mice maintained with 1-2% isoflurane and 95% oxygen to maintain the heart rate at 450±50 beats per minute. Briefly, trans-thoracic echocardiography was conducted using the Vevo F2 Imaging System (#53699-20, FUJIFILM VisualSonics, Inc., USA). The anterior chest hair was removed using Nair Hair-removal cream before sedation. Body-temperature-maintained, prewarmed ultrasound gel was applied to the area underlying the heart after the mice were immobilized on the stage. The parasternal short-axis view, identified by the presence of the papillary muscles, was used to obtain M-mode images for ejection fraction, fractional shortening, left ventricular mass, cardiac output, and other cardiac parameters. The apical four-chamber view was used to get the tissue Doppler and mitral valve pulse-wave Doppler measurements for myocardial tissue and blood flow velocity, respectively. All parameters were measured at least three times, and average measurements were plotted using their means for statistical analysis.

### Histopathology

Histopathology services were performed by the *In Vivo* Animal Core Histology Laboratory within the Unit for Laboratory Animal Medicine at the University of Michigan. Briefly, tissues were fixed in 4% paraformaldehyde (PFA) for 24 hr, then transferred to 70% ethanol. Fixed tissues were embedded in paraffin and sectioned at 4 μm thickness on a rotary microtome (TissueTek VIP5®, Sakura Finetek USA, Inc., Torrance, CA).

Following deparaffinization and hydration with xylene and graded alcohols, formalin-fixed, paraffin-embedded (FFPE) slides were stained with galectin-3 (MAC-2) antibody (sc-81728) as per manufacturer instructions.

### Flow Cytometry

Phenotypic cell analysis of myeloid immune cells in the heart was performed. Briefly, mice were sacrificed, and hearts were rapidly isolated and placed in ice-cold PBS (VWR International). Using a razor blade, the heart was minced manually. After mincing, the tissue was incubated with a mixture of Collagenase type I and type XI (Sigma) and DNase I (Sigma) at 37°C for 30 minutes, with gentle agitation on a rocker. Enzymatic digestion was quenched with cold staining buffer, and the suspension was placed on ice, filtered through a 70 μm cell strainer, then centrifuged at 400 x g for 5 mins at 4°C, followed by debris removal and red blood cell (RBC) lysis steps. The supernatant was discarded, the pellet was resuspended in 1 ml of staining buffer, and the cells were counted. Approximately one million cells were aliquoted per sample for staining.

Cells for flow cytometry were incubated immediately with surface-stain conjugated primary antibodies against RB780conjugated Ly6G (BD Pharmingen), BUV563-conjugated F4/80 (BD), AF488-conjugated LYVE1 (eBioscience), BUV615-conjugated CD11b (BD), APC-CY7-conjugated CD45 (Biolegend), PE-Cy5-conjugated CD64 (Biolegend), BV711-conjugated CD68 (Biolegend), BUV661-conjugated mCCR2 (BD), BV421-conjugated CD115 (Biolegend), BV605-conjugated CD161 (Biolegend), BV570-conjugated CD3 (Biolegend), BUV805-conjugated CD8a (Cytek Biosciences), BUV737-conjugated CD4 (eBioscience) and BUV496-conjugated CD19 (Cytek Biosciences) for 30 minutes on ice. Cells were then washed with staining buffer, permeabilized, and fixed at room temperature for 15 minutes. Cytek Aurora Spectral Analyzer then analyzed samples in the University of Michigan Flow Cytometry Core. Using unstained cells and single fluorescent controls, laser calibration and compensation were performed for all experiments.

#### Data preprocessing

Spectral unmixing and compensation were performed in FlowJo v10.10.0. Events were sequentially gated for intact cells (FSC-A vs SSC-A), singlet discrimination (FSC-A vs FSC-H), viability (Ghost Dye UV450), and CD45+ selection (APC-Cy7). For cardiac tissue, gating thresholds were adjusted to minimize inclusion of autofluorescent cardiomyocytes, as confirmed by comparison with unstained controls. Compensated, scaled values for CD45+ events were exported as CSV files, yielding a combined dataset of 933,463 cells across 15 markers from 16 samples (4 groups × 4 replicates, paired heart and spleen per animal).

#### Manual gating analysis

In parallel, conventional biaxial gating in FlowJo was used to quantify cardiac myeloid populations. CD45+ leukocytes were sequentially gated for non-myeloid markers, CD11b, and Ly6G (neutrophils). CD11b+Ly6G− cells were further assessed for CCR2, F4/80, and CD64. CD45hi/Ly6Glo/F4-80hi cells were identified as macrophages. Neutrophils were defined as CD45hi/CD115lo/Ly6Ghi. Populations were quantified as a percentage of CD45+ cells. Data are presented as mean ± SEM. *P < 0.05.

### RNAseq analyses

Total mRNA was extracted from 20-30 mg of cardiac tissue using TRI Reagent (T9424, Sigma Aldrich, St. Louis, MO, USA) and TissueLyser LT (Qiagen, Hilden, Germany) at 50 Hz. The concentration and purity of mRNA were assessed using a NanoDrop 2000 Spectrophotometer (Thermo Scientific, USA) before synthesizing cDNA from 1000 ng of total RNA from each sample using the Maxima First Strand cDNA Synthesis Kit (#K1671, Thermo Scientific) as per the kit’s protocol on the T100 Thermal Cycler (Applied Biosystems).

#### RNA Sequencing and Differential Gene Expression Analysis

Total RNA was extracted from the heart using the RNeasy Plus Mini Kit (Qiagen), following the manufacturer’s instructions. Novogene performed RNA library preparation and transcriptome sequencing. After quality control, 150 bp paired-end sequencing was performed using the Illumina PE150 platform. Transcriptomic data analysis and visualization were performed using Pluto (https://pluto.bio). Paired-end FASTQ files were processed with the nf-core/rnaseq pipeline (v3.17.0). Briefly, reads were trimmed with Trim Galore, aligned to the Mus musculus GRCm39 reference genome using STAR, and quantified to gene-level counts with RSEM.

Differential gene expression analysis was conducted using the DESeq2 R package, which applies a negative binomial model. Genes with fewer than 3 counts in at least 20% of samples in any group were filtered out before analysis. Log2 fold changes were calculated. Statistical significance was determined using a false discovery rate (FDR)-adjusted p-value threshold of 0.05. Volcano plots display log2 fold changes on the x-axis and -log10 adjusted p-values on the y-axis. Genes with an adjusted p-value ≤ 0.05 and a fold change < –0.6 or > 0.6 were considered significantly altered. Up to 12 of these genes were highlighted on the volcano plot.

#### Gene Set Enrichment Analysis (GSEA)

Gene set enrichment analysis was performed using the fgsea R package and the fgseaMultilevel() function. Genes were ranked by log2 fold change from the DESeq2 output. C2: Canonical pathways - REACTOME gene set collection from the Molecular Signatures Database (MSigDB) was curated using the msigdbr R package. Gene sets were filtered to include only those containing 5-1,000 genes. Normalized enrichment scores (NES) represent the magnitude and direction of enrichment: positive NES values (shown in red) indicate enrichment in the first group, whereas negative NES values (shown in blue) indicate enrichment in the second group. Statistical significance was assessed using p-values.

#### Cell Deconvolution of Bulk RNA-seq Data Using BayesPrism

To perform cellular deconvolution of bulk RNA-seq data, we applied the BayesPrism R package (v2.2.2) in accordance with the authors’ instructions and package vignette (https://github.com/Danko-Lab/BayesPrism). As a reference single-cell dataset, we used publicly available single-cell RNA-seq data from the Gene Expression Omnibus (GSE249412), generated from the same mouse model of HFpEF. Cell types were annotated using canonical markers of major cardiac populations. Because further subdivision into subpopulations was not required for this study, we used identical labels for “cell state” and “cell type.” Lowly expressed genes were removed using the default BayesPrism outlier filtering parameters (outlier.cut = 0.01, outlier.fraction = 0.1). The processed single-cell reference and bulk matrices were then supplied to BayesPrism through the new.prism() function with default settings, and deconvolution was performed using the run.prism() function. Reconstructed expression profiles were subsequently obtained with the get.exp() function.

### In vitro Assessment of the Inflammatory Effects of Soluble uPAR in Bone Marrow–Derived Macrophages

We evaluated the pro-inflammatory effects of uPAR (R&D Systems, Catalog # 531-PA) using murine bone marrow–derived macrophages (BMDMs), as previously published [*Mendoza et al.*, *2022*]. Following differentiation, BMDMs were pre-incubated with 10ng/ml of uPAR for 24 h, after which lipopolysaccharide (LPS, 100 ng/ml and IFN-γ (20 ng/ml) were added for 4 h at 37 °C in a 5% CO₂ incubator. Total RNA was extracted from BMDMs using the RNeasy Plus Mini Kit (Qiagen), following the manufacturer’s instructions.

### Statistical analysis

Data are presented as mean ± SEM. The data were assessed for normality using the Shapiro-Wilk test, and all statistical analyses were performed on log-transformed data when not normally distributed, using GraphPad Prism (Version 10.2.2 [341]). One-way ANOVA with Bonferroni’s post hoc analysis was performed for group comparison. P<0.05 was considered statistically significant.

## Results

### Circulating suPAR overexpression amplifies the cardiometabolic HFpEF phenotype

To dissect how elevated circulating suPAR shapes the HFpEF myocardium, we used the well-validated “two-hit” cardiometabolic mouse model, using high-fat diet (HFD, 60 kcal% fat) plus the constitutive nitric oxide synthase inhibitor Nω-nitro-L-arginine methyl ester (L-NAME), which reproduces the cardinal features of human cardiometabolic HFpEF in C57BL/6 mice (Schiattarella et al., 2019), in the murine suPAR-transgenic (suPAR-Tg) strain. These mice have previously been shown to circulate suPAR at concentrations approximating those observed in cardiovascular and kidney disease and to accelerate atherosclerosis by priming circulating monocytes (Hayek et al., 2017; Hindy et al., 2022). Eight- to ten-week-old male wild-type (WT) and suPAR-Tg littermates were assigned to one of three groups: LFD-WT, HFD+L-NAME-WT, or HFD+L-NAME-suPAR-Tg, and maintained on the assigned regimen for 15 weeks (**Figure 1A**). The choice of this model is motivated by the consistent epidemiologic observation that suPAR is independently associated with adverse HFpEF outcomes beyond natriuretic peptides (Hayek et al., 2023; Hutten et al., 2025), yet whether suPAR is causally involved or merely a passive marker of inflammatory burden has remained unresolved (Reiser et al, 2026).

**Figure 1.**
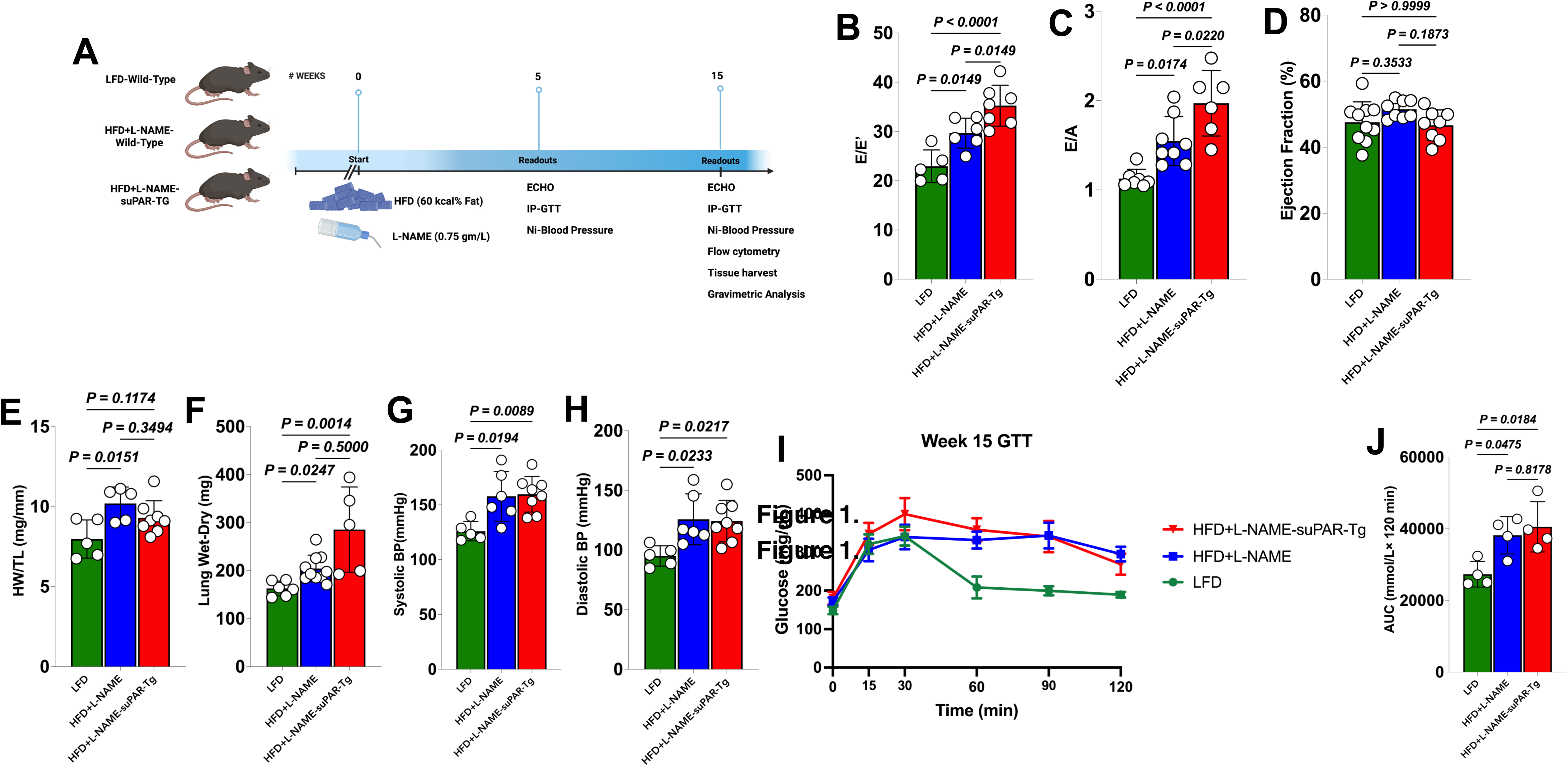
suPAR worsens diastolic dysfunction, increases pulmonary congestion, has a modest impact on blood pressure, and impairs glucose tolerance in a diet-induced cardiometabolic syndrome regimen. (A) Schematic of the experimental protocol. Wild-type (WT) and suPAR-transgenic (suPAR-Tg) mice were assigned to one of three groups — LFD-WT, HFD+L-NAME-WT, or HFD+L-NAME-suPAR-Tg — and fed either a low-fat diet (LFD) or a high-fat diet (HFD, 60 kcal% fat) combined with L-NAME (0.75 g/L) in the drinking water for 15 weeks. Baseline echocardiography (echo), body weight, and blood pressure were obtained prior to randomization. Interim readouts at week 5 included echo, IP-GTT, and non-invasive blood pressure. Endpoint measurements at week 15 included echo, IP-GTT, non-invasive blood pressure, flow cytometry, tissue harvest with histology, and gravimetric analyses. (B) Ratio of peak early diastolic mitral inflow velocity (E) to peak early diastolic mitral annular velocity (e’) (E/e’ ratio). (C) Ratio of peak early (E) to late (A) diastolic mitral inflow velocity (E/A ratio), measured by pulsed-wave Doppler. (D) Left ventricular ejection fraction (EF, %). (E) Gravimetric heart weight normalized to tibia length (HW/TL, mg/mm). (F) Lung water content, assessed by the difference between wet and dry lung weight (mg). (G–H) Endpoint (week 15) systolic blood pressure (G) and diastolic blood pressure (H). (I) Blood glucose during an intraperitoneal glucose tolerance test (IP-GTT) performed at the study endpoint (week 15). (J) Area under the curve (AUC) quantification for the GTT is shown in (I). Each symbol represents an individual animal; bars represent mean ± SEM. Statistical significance was determined by one-way or two-way ANOVA, as appropriate, with Bonferroni post hoc tests. LFD indicates low-fat diet; HFD, high-fat diet; L-NAME, Nω-nitro-L-arginine methyl ester; echo, echocardiography; IP-GTT, intraperitoneal glucose tolerance test; HW/TL, heart weight/tibia length; BP, blood pressure; AUC, area under the curve.

Mitral inflow and tissue Doppler interrogation at week 15 revealed a stepwise worsening of diastolic function. The E/e′ ratio, a non-invasive surrogate of left ventricular filling pressure, was higher in HFD+L-NAME-WT than in LFD-WT (P = 0.0149) and rose further in HFD+L-NAME-suPAR-Tg (P = 0.0149 vs HFD+L-NAME-WT; P < 0.0001 vs LFD-WT) (**Figure 1B**). The E/A ratio followed the same trajectory, with the highest values observed in HFD+L-NAME-suPAR-Tg (P = 0.0220 vs HFD+L-NAME-WT; P < 0.0001 vs LFD-WT) (**Figure 1C**). LV ejection fraction remained preserved and indistinguishable across groups (**Figure 1D**), confirming an HFpEF phenotype rather than a drift toward systolic failure.

#### Gravimetric endpoints paralleled the functional data

Heart weight normalized to tibia length (HW/TL) increased in HFD+L-NAME-WT compared with LFD-WT (P = 0.0151) but did not increase further in HFD+L-NAME-suPAR-Tg (P = 0.3494) (**Figure 1E**), indicating that suPAR does not augment pressure-load-driven hypertrophy beyond what the cardiometabolic stimulus already produces. Lung wet-to-dry weight was, by contrast, markedly elevated only in HFD+L-NAME-suPAR-Tg (P = 0.0014 vs LFD-WT; P = 0.0247 vs HFD+L-NAME-WT) (**Figure 1F**). Systolic and diastolic blood pressures were both higher in the HFD+L-NAME groups compared with LFD-WT (systolic, P ≤ 0.0194; diastolic, P ≤ 0.0233) but did not differ between the two HFD+L-NAME groups (**Figure 1G–H**), confirming that the additional impairment observed in suPAR-Tg animals is not attributable to differential afterload. Glucose handling was impaired by HFD+L-NAME in both WT and suPAR-Tg animals during the IP-GTT (**Figure 1I**) and a significantly increased area under the curve (P = 0.0184) (**Figure 1J**), consistent with prior evidence that suPAR levels track with incident type 2 diabetes and insulin resistance in human cohorts (Eugen-Olsen et al., 2010).

Together, these findings establish that systemic suPAR elevation in the cardiometabolic two-hit model produces a more severe HFpEF phenotype, with greater diastolic dysfunction and pulmonary congestion, albeit without altering blood pressure, suggesting a mechanism operating downstream of the canonical hemodynamic stimulus.

### suPAR drives a cardiac transcriptomic shift toward immune activation and metabolic suppression in HFpEF

To define the molecular basis of the worsened phenotype, we performed bulk RNA sequencing on left ventricular tissue from HFD+L-NAME-WT and HFD+L-NAME-suPAR-Tg mice at the 15-week endpoint. Gene Set Enrichment Analysis (GSEA) using the C2/REACTOME and Gene Ontology collections revealed a coordinated bidirectional shift (**Figure 2A**): pathways enriched in suPAR-Tg hearts centered on leukocyte aggregation, T-cell chemotaxis and migration, interleukin-1β (IL-1β) production, and medium-chain fatty acid biosynthesis, while pathways enriched in WT hearts centered on oxidative phosphorylation (OXPHOS), the mitochondrial respiratory chain and NADH dehydrogenase complex assembly, mitochondrial translation, long-chain fatty acid transport, arachidonic acid and icosanoid secretion, and PPAR signaling. This combination of suppressed mitochondrial metabolism and amplified innate immunity recapitulates the metabolic-inflammatory signature central to the modern HFpEF paradigm (Paulus & Tschöpe, 2013; Paulus & Zile, 2021; Schiattarella et al., 2021).

**Figure 2.**
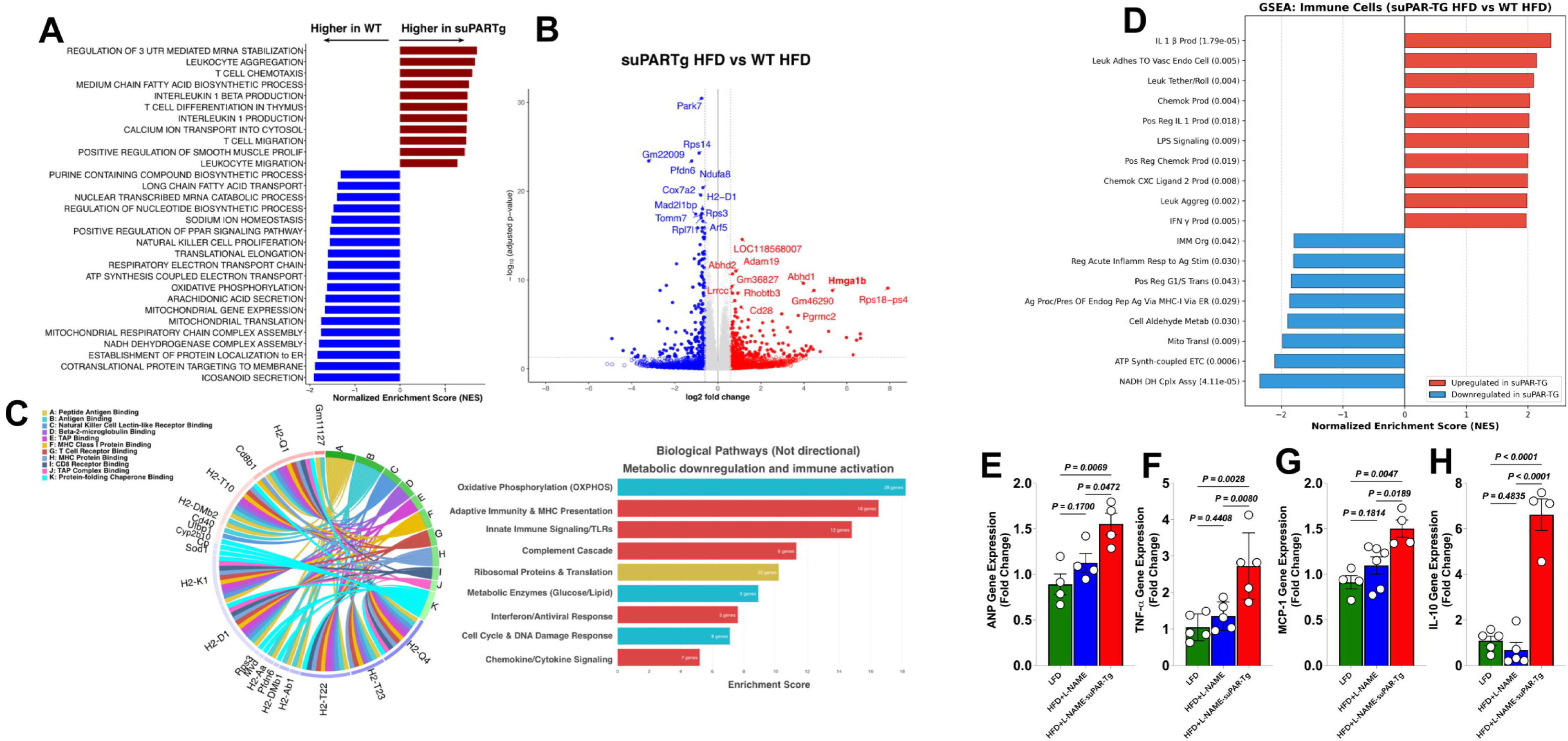
suPAR increases HFpEF-induced cardiac transcriptomic alterations, particularly in immune cells. (A) Gene Set Enrichment Analysis (GSEA) comparing pathway activity in cardiac tissue between HFD+L-NAME-WT and HFD+L-NAME-suPAR-Tg mice. Bars represent the Normalized Enrichment Score (NES). Red indicates pathways enriched in the suPAR-Tg group, while blue indicates pathways depleted (enriched in WT). The data suggest that suPAR induces pathways involved in inflammation and immune cell activation while suppressing mitochondrial oxidative phosphorylation, fatty acid transport, and icosanoid metabolism. (B) Volcano plot showing differentially expressed genes (DEGs) in cardiac tissue comparing HFD+L-NAME-suPAR-Tg vs. HFD+L-NAME-WT. The x-axis represents log2 fold change and the y-axis represents –log10 adjusted P value. Significantly upregulated genes are shown in red and downregulated genes in blue (significance threshold: adjusted P < 0.05). Key labeled genes represent those involved in fatty acid and glucose metabolism and in inflammatory pathways, including upregulated Hmga1b, Pgrmc2, Cd28, Adam19, and Abhd1/2, and downregulated Park7, Ndufa8, Cox7a2, Tomm7, and H2-D1. (C) Pathway enrichment showing immune/inflammation gene signatures upregulated in suPAR-Tg hearts. The chord diagram (left) links individual MHC and immune-related transcripts (H2-K1, H2-D1, H2-Q1, H2-Q4, H2-T10/T22/T23, H2-Aa, H2-Ab1, H2-DMb1/2, Cd8b1, Cd40, Klrk1, Ulbp1) to molecular function categories (A–K: peptide antigen binding, antigen binding, NK cell lectin-like receptor binding, β2-microglobulin binding, TAP binding, MHC class I protein binding, T-cell receptor binding, MHC protein binding, CD8 receptor binding, TAP complex binding, and protein-folding chaperone binding). The bar chart (right) summarizes non-directional pathway enrichment spanning oxidative phosphorylation, ribosomal proteins and translation, adaptive immunity and MHC presentation, innate immune signaling/TLRs, complement cascade, metabolic enzymes (glucose/lipid), interferon/antiviral response, cell cycle and DNA damage response, and chemokine/cytokine signaling. (D) GSEA of pathways altered within the cardiac immune cell populations (derived from inferred pathway activity data), comparing HFD+L-NAME-suPAR-Tg vs. HFD+L-NAME-WT. Red bars indicate enrichment in suPAR-Tg (IL-1β production, leukocyte adhesion to vascular endothelium and tethering/rolling, chemokine production including CXCL2, LPS signaling, leukocyte aggregation, and IFN-γ production); blue bars indicate depletion (NADH dehydrogenase complex assembly, ATP synthesis–coupled electron transport, mitochondrial translation, cellular aldehyde metabolism, MHC-I antigen processing via ER, and regulation of acute inflammatory response). Adjusted P values are shown in parentheses. (E–H) Cardiac gene expression by qPCR shows upregulated heart failure and inflammatory markers in suPAR-Tg hearts: (E) ANP (Nppa), (F) TNF-α (Tnf), (G) MCP-1 (Ccl2), and (H) IL-10 (Il10), measured in mice on low-fat diet (LFD, green), HFD+L-NAME (blue), or HFD+L-NAME-suPAR-Tg (red). Each symbol represents an individual animal (n = 4–6 per group). Data are presented as mean ± SEM. Statistical significance was determined by one-way or two-way ANOVA, as appropriate, with Bonferroni post hoc tests.

Differential expression analysis (DESeq2, adjusted P < 0.05, |log₂ fold change| ≥ 0.585) identified hundreds of transcripts altered by suPAR overexpression (**Figure 2B**). Upregulated genes included transcripts implicated in cardiac stress and inflammation (Hmga1b, Pgrmc2, Cd28, Adam19) and in lipid metabolism (Abhd1, Abhd2, Rhobtb3). Hmga1, an architectural transcription factor that aggravates LPS-induced and diabetic cardiomyopathy (Wu et al., 2020; Wu et al., 2020 Cell Death Dis), and Pgrmc2, a membrane progesterone receptor recently established as a critical regulator of myocardial pressure-volume responses to stress (Thomas et al., 2025), suggest activation of stress-response and remodeling programs in suPAR-exposed myocardium. Downregulated genes were enriched for OXPHOS and mitochondrial translation components (Ndufa8, Cox7a2, Tomm7), the redox sensor Park7/DJ-1, and class I MHC transcripts (H2-D1), consistent with the suppressed mitochondrial program identified by GSEA.

To localize these transcriptional changes to specific cellular compartments, we performed in silico cellular deconvolution of the bulk RNA-seq data using BayesPrism (Chu et al., 2022) with a published single-cell reference derived from an analogous HFpEF mouse model (Tucker et al., 2024). Non-directional pathway enrichment across the deconvolved compartments confirmed a coordinated reorganization (**Figure 2C**): the largest gene sets affected were OXPHOS (28 genes), ribosomal proteins and translation (32 genes), adaptive immunity and MHC presentation (16 genes), innate immune signaling/TLRs (12 genes), and chemokine/cytokine signaling (7 genes). A chord diagram of MHC-related transcripts (H2-K1, H2-D1, H2-Q1/Q4, H2-T10/T22/T23, H2-Aa, H2-Ab1, H2-DMb1/2) and co-receptors (Cd8b1, Cd40, Klrk1, Ulbp1) mapped these signals onto antigen processing, MHC class I/II binding, T-cell receptor binding, β2-microglobulin binding, and TAP-complex binding categories, indicating that suPAR remodels not only the innate but also the antigen-presentation machinery of the HFpEF heart.

When GSEA was restricted to deconvolved immune cell populations, the inflammatory signature was augmented (**Figure 2D**): suPAR-Tg hearts showed strong enrichment for IL-1β production (adjusted P = 1.79 × 10⁻⁵), leukocyte adhesion to vascular endothelium and tethering/rolling, CXCL2 production, LPS signaling, leukocyte aggregation, and IFN-γ production, alongside depletion of NADH dehydrogenase complex assembly (adjusted P = 4.11 × 10⁻⁵), ATP synthesis-coupled electron transport, mitochondrial translation, and MHC-I antigen processing via the endoplasmic reticulum.

Quantitative PCR in independent cohorts validated this signature at the gene level (**Figure 2E–H**): atrial natriuretic peptide (Nppa, P = 0.0069), TNF-α (Tnf, P = 0.0028), MCP-1/Ccl2 (P = 0.0047), and IL-10 (Il10, P < 0.0001) were all increased in HFD+L-NAME-suPAR-Tg hearts. The robust IL-10 induction is mechanistically informative, as macrophage-derived IL-10 in the failing heart actively promotes fibroblast activation, collagen deposition, and diastolic dysfunction (Tucker et al., 2024; Raman et al., 2025), and its upregulation here therefore reflects pathologic, not counter-regulatory, macrophage activation.

### suPAR increases cardiac and systemic myeloid infiltration in HFpEF

We performed manual gating of the spectral flow cytometry on enzymatically digested cardiac tissue, spleen, and peripheral blood mononuclear cells (PBMCs) at 15 weeks (**Figure 3**). In the heart, total myeloid cells (CD45⁺GR-1⁺) increased stepwise across groups, reaching the highest values in HFD+L-NAME-suPAR-Tg (P = 0.0004 vs LFD-WT; P = 0.0123 vs HFD+L-NAME-WT) (**Figure 3A**). Neutrophils (CD45⁺CD11b⁺Ly6G⁺) were markedly elevated in HFD+L-NAME-suPAR-Tg (P = 0.0006 vs LFD-WT; P = 0.0133 vs HFD+L-NAME-WT) (**Figure 3B**).

**Figure 3.**
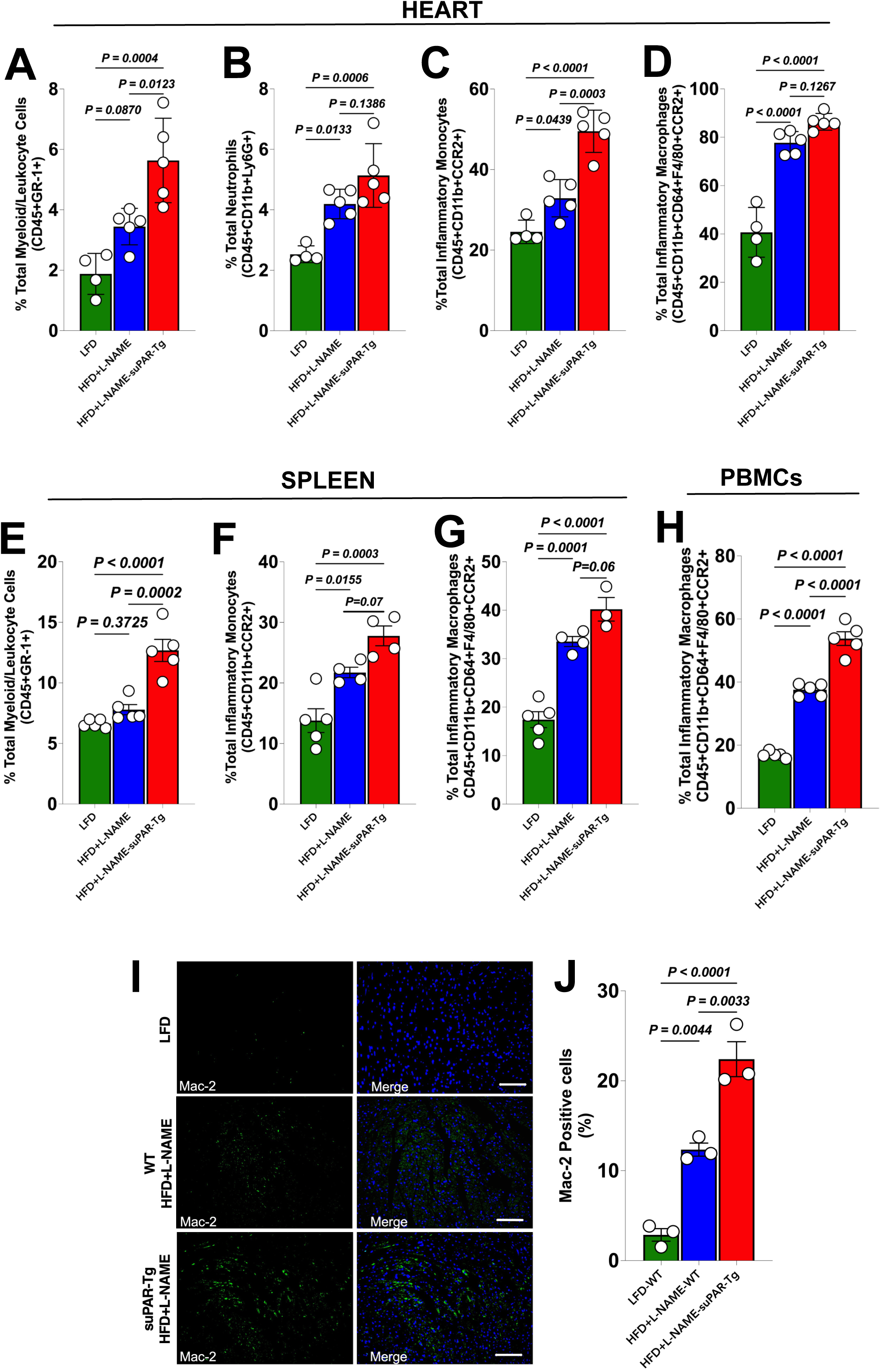
suPAR increases HFpEF-induced cardiac immune-cell infiltration and induces systemic inflammation. (A–D) Flow cytometric quantification of myeloid populations in cardiac tissue. (A) Total myeloid/leukocyte cells (CD45⁺GR-1⁺) are increased in HFD+L-NAME-suPAR-Tg hearts compared with both HFD+L-NAME-WT and LFD-WT. (B) Neutrophils (CD45⁺CD11b⁺Ly6G⁺) are markedly elevated in HFD+L-NAME-suPAR-Tg hearts. (C) Inflammatory monocytes (CD45⁺CD11b⁺CCR2⁺) and (D) inflammatory macrophages (CD45⁺CD11b⁺CD64⁺F4/80⁺CCR2⁺) are increased in HFD+L-NAME-suPAR-Tg hearts. (E–G) Splenic immune cell populations show a parallel pattern of expansion: (E) total myeloid/leukocyte cells (CD45⁺GR-1⁺), (F) inflammatory monocytes (CD45⁺CD11b⁺CCR2⁺), and (G) inflammatory macrophages (CD45⁺CD11b⁺CD64⁺F4/80⁺CCR2⁺). (H) Inflammatory macrophages (CD45⁺CD11b⁺CD64⁺F4/80⁺CCR2⁺) are increased in HFD+L-NAME-suPAR-Tg peripheral blood mononuclear cells (PBMCs). (I–J) Representative immunofluorescence images (I) and quantification (J) demonstrate significantly higher Mac-2⁺ macrophage density in suPAR-Tg hearts compared with HFD+L-NAME-WT and LFD-WT. Mac-2, green; DAPI nuclear counterstain, blue. Scale bars, [confirm length, µm]. Bars represent mean ± SEM; each symbol denotes an individual mouse. Statistical significance was determined by one-way ANOVA with Dunnett post hoc multiple-comparisons test.

Inflammatory bone marrow-derived monocytes (CD45⁺CD11b⁺CCR2⁺) and inflammatory macrophages (CD45⁺CD11b⁺CD64⁺F4/80⁺CCR2⁺) were similarly expanded (monocytes, P < 0.0001 vs LFD-WT; P = 0.0003 vs HFD+L-NAME-WT; macrophages, P < 0.0001 vs LFD-WT) (**Figure 3C–D**). The selective expansion of the CCR2⁺ inflammatory subsets — recruited from bone marrow and spleen, distinct from embryonic-derived resident macrophages, and required for adverse remodeling in pressure-overload, dyslipidemic, and early HFpEF models (Patel et al., 2018; Tucker et al., 2024; Raman et al., 2025) — identifies monocyte recruitment, rather than resident proliferation, as the dominant mechanism.

The systemic immune compartment was similarly affected. Splenic myeloid/leukocyte cells (CD45⁺GR-1⁺), inflammatory monocytes, and inflammatory macrophages were each increased in HFD+L-NAME-suPAR-Tg (P < 0.0001 vs LFD-WT for all comparisons) (**Figure 3E–G**), suggesting active extramedullary myelopoiesis. CCR2⁺ inflammatory macrophages were also significantly more abundant in PBMCs (P < 0.0001 vs both control groups) (**Figure 3H**), demonstrating that the inflammatory state extends beyond the heart. These findings extend prior observations that suPAR-Tg mice circulate a primed monocyte pool with enhanced chemotactic capacity in the setting of atherosclerosis (Hindy et al., 2022), and identify the cardiometabolic HFpEF context as a second disease in which this primed myeloid pool produces pathology.

To corroborate the flow cytometric findings in situ, we performed immunofluorescence for galectin-3 (Mac-2), a marker of activated tissue macrophages with established prognostic value in heart failure (de Boer et al., 2009). Mac-2⁺ cell density was substantially higher in HFD+L-NAME-suPAR-Tg hearts than in HFD+L-NAME-WT (P = 0.0033) or LFD-WT (P < 0.0001) (**Figure 3I–J**), spatially confirming the recruited macrophage population identified by flow cytometry. Collectively, these data establish that suPAR amplifies myocardial inflammation in HFpEF by promoting both cardiac and systemic expansion of CCR2⁺ inflammatory myeloid populations.

### suPAR primes bone marrow–derived macrophages for amplified inflammatory responses

A central unresolved question is whether suPAR acts directly on macrophages to drive the inflammatory phenotype or whether the cardiac findings reflect indirect effects mediated by hemodynamic, metabolic, or endothelial changes. While suPAR signals through the αvβ3 integrin in podocytes (Hayek et al., 2017) and primes circulating monocytes for enhanced chemotaxis in atherosclerosis (Hindy et al., 2022), direct evidence that recombinant suPAR potentiates macrophage cytokine responses has not been reported. To address this gap, we differentiated bone marrow–derived macrophages (BMDMs) from naïve C57BL/6 mice and pretreated them with recombinant mouse suPAR (10 ng/mL) or vehicle for 24 hours, followed by stimulation with LPS (100 ng/mL) and IFN-γ (20 ng/mL) for 4 hours (**Figure 4A**), a dual stimulus that mimics the TLR4-driven, IFN-γ-amplified state of inflammatory macrophages.

**Figure 4.**
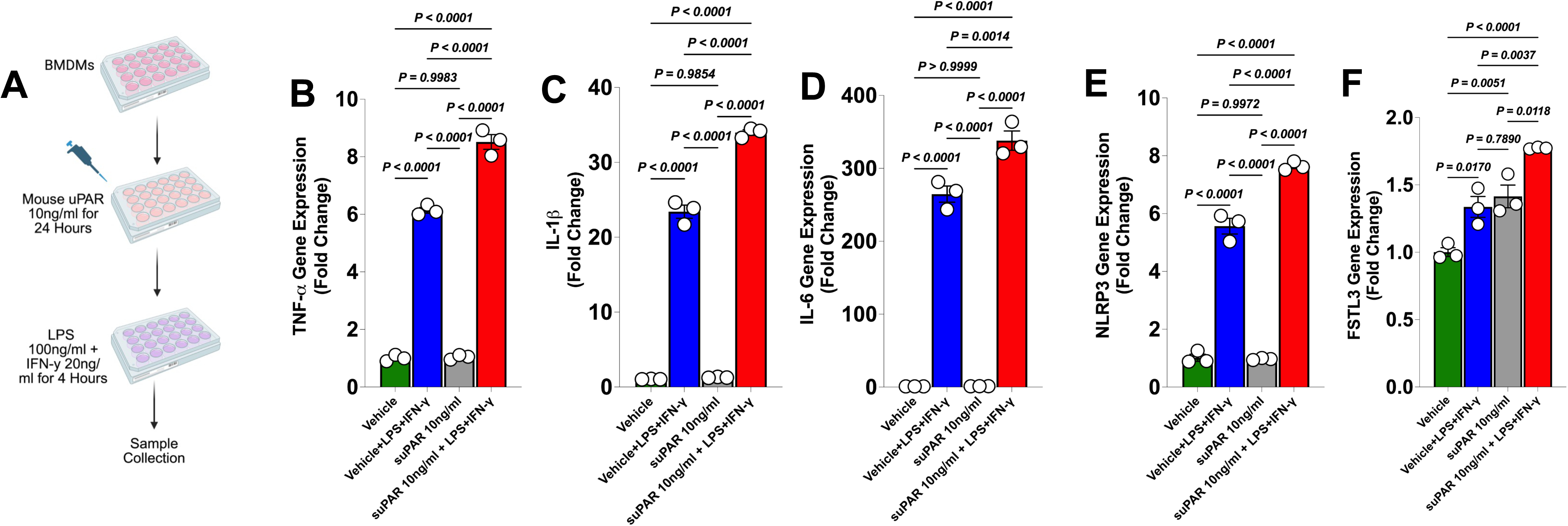
suPAR increases inflammatory gene expression in bone marrow-derived macrophages (BMDMs) upon LPS and IFN-γ induction. (A) Schematic of the in vitro experimental design. BMDMs were pretreated with recombinant mouse suPAR (10 ng/mL) or vehicle for 24 hours, followed by stimulation with LPS (100 ng/mL) and IFN-γ (20 ng/mL) for 4 hours prior to sample collection. (B–F) Gene expression analysis by qPCR for (B) TNF-α, (C) IL-1β, (D) IL-6, (E) NLRP3, and (F) FSTL3, expressed as fold change relative to vehicle. Four conditions are shown in each panel: vehicle (green), vehicle + LPS+IFN-γ (blue), suPAR 10 ng/mL alone (gray), and suPAR 10 ng/mL + LPS+IFN-γ (red). suPAR pretreatment alone did not induce TNF-α, IL-1β, IL-6, or NLRP3 expression but markedly potentiated their induction by LPS+IFN-γ; FSTL3 was modestly elevated by suPAR alone and further amplified by combined stimulation. Each symbol represents an individual replicate; bars represent mean ± SEM. Statistical significance was determined by one-way ANOVA with Bonferroni post hoc tests for multiple comparisons.

suPAR pretreatment alone did not significantly induce Tnf, Il1b, Il6, or Nlrp3 compared with vehicle (P ≥ 0.99 for all comparisons), establishing that soluble uPAR is not a stand-alone inflammatory ligand at this concentration (**Figure 4B–E**). However, when suPAR pretreatment was combined with LPS+IFN-γ, gene expression of all four inflammatory mediators was further amplified compared with LPS+IFN-γ alone: TNF-α (P < 0.0001), IL-1β (P < 0.0001), IL-6 (P = 0.0014), and NLRP3 (P < 0.0001). The induction of NLRP3 is mechanistically significant: NLRP3 inflammasome activation, engaged by mitochondrial damage-associated molecular patterns released from the metabolically stressed HFpEF heart, is an emerging node linking metabolic stress to diastolic dysfunction (Paulus & Zile, 2021; Schiattarella et al., 2021), and our data identify suPAR as an upstream amplifier of this pathway. Fstl3, encoding follistatin-like 3, a TGF-β superfamily antagonist implicated in cardiac hypertrophy and fibrosis (Panse et al., 2012), was modestly increased by suPAR alone (P = 0.0170) and further amplified by combined stimulation (P < 0.0001), suggesting that suPAR engages both a direct hypertrophic-remodeling program and an inflammasome-priming program in macrophages (**Figure 4F**).

These *in vitro* data identify suPAR as a macrophage-priming factor rather than a stand-alone inflammatory ligand: it lowers the threshold for, and amplifies the magnitude of, the canonical TLR4/IFN-γ inflammatory response. This mechanism provides a parsimonious explanation for the in vivo findings in HFD+L-NAME-treated suPAR-Tg mice, where TLR ligands (oxidized lipids, damage-associated molecular patterns) and pro-inflammatory cytokines are abundant, sustained suPAR exposure converts a moderate cardiometabolic insult into a more aggressive immune-driven HFpEF phenotype, and provides the first direct molecular evidence supporting the prognostic association between circulating suPAR and HFpEF outcomes observed in clinical cohorts (Hayek et al., 2023; Hutten et al., 2025).

## Discussion

This study identifies suPAR, the soluble form of the urokinase plasminogen activator receptor, as a direct, mechanistic amplifier of HFpEF in a well-characterized cardiometabolic mouse model. Three lines of evidence support this conclusion. First, transgenic overexpression of suPAR in mice subjected to the high-fat diet plus L-NAME two-hit regimen (Schiattarella et al., 2019) produced a more severe HFpEF phenotype than the cardiometabolic insult alone, with greater elevation in E/e′ and greater pulmonary congestion. Second, the cardiac transcriptome of suPAR-Tg HFpEF mice showed a coordinated bidirectional shift, with suppression of mitochondrial oxidative phosphorylation and amplification of innate and adaptive immune pathways, accompanied by demonstrable expansion of CCR2⁺ inflammatory monocytes and macrophages in the heart, spleen, and peripheral blood. Third, recombinant suPAR primed bone marrow–derived macrophages for amplified TNF-α, IL-1β, IL-6, and NLRP3 induction upon LPS+IFN-γ stimulation, providing the first direct in vitro evidence, to our knowledge, that soluble uPAR sensitizes macrophages to canonical inflammatory stimuli. Together, these findings transform suPAR from a passive biomarker of inflammation into a sufficient upstream amplifier of the inflammatory cascade that defines HFpEF.

The data presented here are best explained by a model in which circulating suPAR engages the αvβ3 integrin on monocytes and tissue macrophages, lowering the activation threshold for canonical pattern-recognition receptor signaling. The αvβ3 mechanism is well-established in podocytes, where the suPAR-APOL1-αvβ3 tripartite complex drives focal segmental glomerulosclerosis (Wei et al., 2008; Hayek et al., 2017), and αvβ3 ligation alone potentiates NF-κB-dependent TNF-α and IL-1β responses in human monocytes by an order of magnitude.

The macrophage-priming pattern we observed, demonstrating no induction by suPAR alone but markedly amplifying TNF-α, IL-1β, IL-6, and NLRP3 upon LPS+IFN-γ co-stimulation, mirrors this integrin-priming phenomenon and extends it from kidney to cardiac biology. It also aligns with the in vivo phenotype previously described in suPAR-Tg mice, in which circulating monocytes are preprogrammed for enhanced chemotaxis and entry into atherosclerotic lesions (Hindy et al., 2022).

The NLRP3 induction is mechanistically informative. NLRP3 inflammasome activation occupies a central position in the contemporary HFpEF paradigm, engaged by mitochondrial damage-associated molecular patterns released from metabolically stressed cardiomyocytes (Paulus & Zile, 2021; Schiattarella et al., 2021). Our data place suPAR upstream of this node: by priming macrophages for amplified NLRP3 expression, suPAR enables a stronger inflammasome response to the same metabolic insult. This positioning is consistent with the in vivo finding that mitochondrial OXPHOS and electron transport pathways were among the most strongly suppressed gene sets in suPAR-Tg HFpEF hearts (adjusted P = 4.11 × 10⁻⁵ for NADH dehydrogenase complex assembly), suggesting a metabolic substrate that the primed macrophages then translate into amplified inflammation.

The upregulation of Il10 in suPAR-Tg hearts may appear paradoxical given the inflammatory phenotype, but the contemporary cardiac macrophage literature has reframed this signal. Macrophage-derived IL-10 in the failing heart actively promotes fibroblast activation, collagen deposition, and diastolic stiffening rather than serving a counter-regulatory role (Tucker et al., 2024; Raman et al., 2025). Our finding, therefore, reflects pathologic activation of the recruited CCR2⁺ macrophage compartment, not a homeostatic restraint of inflammation, and provides additional mechanistic support for the diastolic phenotype observed in Figure 1.

Evidence that suPAR is associated with HFpEF outcomes has steadily accumulated over the past decade. In broader heart failure cohorts, suPAR independently predicts mortality and hospitalization beyond natriuretic peptides (Hayek et al., 2023). In the TOPCAT North American cohort, baseline suPAR in the highest tertile conferred more than a twofold increase in the risk of cardiovascular death, cardiac arrest, or heart failure hospitalization, with risk discrimination improved by 0.123 C-statistic points beyond age, sex, race, and natriuretic peptides (Hutten et al., 2025). Crucially, spironolactone, the only mineralocorticoid receptor antagonist with potential HFpEF activity, did not lower suPAR levels, and its therapeutic effect was not modified by baseline suPAR. The most parsimonious interpretation of those observations, combined with the present mechanistic findings, is that suPAR identifies an inflammatory phenotype that is upstream of and orthogonal to mineralocorticoid receptor blockade.

Our findings are also consistent with, and mechanistically extend, the emerging recognition that CCR2⁺ monocyte-derived macrophages, rather than embryonically derived resident macrophages, drive HFpEF remodeling. Patel et al. (2018) established that genetic ablation of CCR2⁺ monocyte-derived macrophages attenuate pressure-overload remodeling, and Tucker et al. (2024) and Raman et al. (2025) extended this paradigm to dyslipidemia- and pressure-volume-driven early HFpEF, respectively. We add to this body of work by identifying an upstream circulating signal, suPAR, that selectively expands and primes the same CCR2⁺ subset. This positions suPAR as a potential therapeutic entry point into an immune mechanism that has become increasingly difficult to target through downstream cytokine blockade.

### Translational implications

Three decades of anti-inflammatory clinical trials in cardiovascular disease have produced mixed results. CANTOS demonstrated that IL-1β blockade with canakinumab reduces atherothrombotic events (Ridker et al., 2017) and improves HF hospitalization in patients with elevated hsCRP (Everett et al., 2018) but failed to gain regulatory approval for cardiovascular indications. Ziltivekimab, an IL-6 ligand antibody, lowers hsCRP by >85% in patients with HF and elevated CRP and is now in the Phase 3 ZEUS trial (Ridker et al., 2021), but mortality outcomes are pending. The four available therapies with HFpEF mortality or morbidity signals, spironolactone, sacubitril-valsartan, empagliflozin (Anker et al., 2021), and dapagliflozin (Solomon et al., 2022), were not designed around inflammation, and the magnitude of benefit, while real, remains modest. The persistent gap suggests that broad downstream cytokine blockade may not capture the disease’s upstream inflammatory architecture.

Our findings provide a mechanistic argument for upstream targeting. A humanized anti-suPAR monoclonal antibody (WAL0921; Walden Biosciences, 2025) is currently in a Phase 2 basket study for chronic kidney diseases, with interim readouts expected in 2026. Given the substantial epidemiologic overlap between cardiorenal syndrome and HFpEF, and the shared αvβ3-integrin mechanism in both organs (Reiser, 2026), the suPAR-Tg HFpEF model described here provides a well-defined preclinical platform for evaluating the efficacy of anti-suPAR antibodies in HFpEF prior to dedicated cardiovascular trials. Our findings also identify NLRP3 as a downstream node that could be co-targeted with anti-suPAR therapy, a combination strategy that may capture both the upstream priming signal and the terminal inflammasome amplifier.

### Study limitations

The study limitations warrant explicit acknowledgment. First, only male mice were studied because female mice are less susceptible to the HFD+L-NAME phenotype [*Tong et al.*, *2019*]; however, suPAR levels are sexually dimorphic in humans, and the sex-specific consequences of suPAR overexpression in HFpEF will require dedicated study. Second, the transgenic model produces persistent and supraphysiologic suPAR elevation; whether transient or intermediate elevations, closer to those observed in cardiometabolic patients, produce comparable cardiac phenotypes remains to be determined. Third, bone marrow–derived macrophages are not identical to cardiac tissue macrophages, and the priming effect we documented may differ quantitatively or qualitatively between resident and recruited cardiac populations; ongoing work with single-cell preparations will address this gap. Fourth, the bulk RNA-seq deconvolution provides cell-type-level resolution but is necessarily inferential; direct scRNA-seq in suPAR-Tg HFpEF hearts will be required to definitively localize the transcriptional changes and to test whether the suPAR effect operates predominantly through monocyte recruitment, resident macrophage priming, or both. Fifth, the *in vivo* model is gain-of-function: sustained transgenic suPAR overexpression establishes that elevated circulating suPAR is sufficient to worsen the cardiometabolic HFpEF phenotype and to expand and prime the CCR2⁺ macrophage compartment, but it does not establish necessity; loss-of-function studies using suPAR neutralization and conditional deletion are ongoing to test whether suPAR is required for the inflammatory remodeling described here.

## Conclusions

Our findings establish that elevated soluble urokinase plasminogen activator receptor is sufficient to act as an upstream amplifier of HFpEF in a cardiometabolic mouse model, acting by priming bone marrow–derived macrophages to augment inflammatory responses. The data link two decades of epidemiologic association between suPAR and adverse heart failure outcomes to a mechanistic and therapeutic hypothesis and identify an immediately tractable translational opportunity in the form of clinical-stage anti-suPAR antibodies. Whether neutralization of circulating suPAR can attenuate established HFpEF and which patients are most likely to benefit are the questions this work most directly invites.

## Acknowledgements.

A.A.L was supported by VA Merit (I01CX002684-01), NIH R61/R33 (1R61HL177474-01), and the Mathers Foundation grant. Additional funds from Lexicon supported this project.

An AHA postdoctoral fellowship supports A.A.L., R.C., and R.L.

S.N.G. was supported by VA MERIT grant 1I01CX002560 and the Taubman Medical Research Institute (Wolfe Scholarship)

S.S.H. is supported by NIH grant 1R01HL153384-01.

## Data sharing statement

The data underlying this article will be shared on reasonable request to the corresponding author.

